# Early life adversity shapes life history trade-offs between growth and reproduction in free-ranging rhesus macaques

**DOI:** 10.1101/2025.09.11.675677

**Authors:** Rachel M. Petersen, Sam K. Patterson, Anja Widdig, Cassandra M. Turcotte, Susan C. Antón, Scott A. Williams, Ashly N. Romero, Samuel E. Bauman Surratt, Angelina V. Ruiz Lambides, Cayo Biobank Research Unit, Michael J. Montague, Noah Snyder-Mackler, Lauren J.N. Brent, James P. Higham, Amanda J. Lea

**Author notes:** These authors contributed equally (Rachel M. Petersen, Sam K. Patterson).

## Abstract

Life history theory predicts that organisms allocate resources across physiological processes to maximize fitness. Under this framework, early life adversity (ELA)—which often limits energetic capital—could shape investment in growth and reproduction, as well as trade-offs between them, ultimately contributing to variation in evolutionary fitness. Using long-term demographic, behavioral, and physiological data for 2,100 females from a non-human primate population, we tested whether naturally-occurring ELA influences investment in the competing physiological demands of growth and reproduction. By analyzing ELA, growth, and reproduction in the same individuals, we also assessed whether adversity intensifies trade-offs between life history domains. We found that ELA influenced life history patterns, and was associated with modified growth, delayed reproductive maturity, and small adult body size. Different types of ELA sometimes had distinct reproductive outcomes—e.g., large group size was linked to faster reproductive rates, while low maternal rank predicted slower ones. Adversity also amplified trade-offs between growth and reproduction: small body size was a stronger predictor of delayed and reduced reproductive output in females exposed to ELA, compared to those not exposed. Finally, we examined how traits modified by ELA related to lifetime reproductive success. Across the population, starting reproduction earlier and maintaining a moderate reproductive rate conferred the greatest number of offspring surviving to reproductive maturity. These findings suggest that ELA impacts key life history traits as well as relationships between them, and can constrain individuals from adopting the most optimal reproductive strategy.

**Significance Statement:** Early life adversity (ELA) can have lasting effects on evolutionary fitness (e.g., the number of surviving offspring an animal produces); however, the paths connecting ELA to fitness—for example by influencing growth, reproductive timing or rate, or trade-offs between these processes—remain unclear. Leveraging long-term behavioral, physiological, and demographic data from 2,100 female rhesus macaques, we found that ELA-exposed females exhibited growth and reproductive schedules associated with less-optimal lifetime fitness outcomes. Further, ELA intensified trade-offs between growth and reproduction, suggesting that affected individuals face steeper energetic constraints. Our findings highlight the long-lasting impacts of ELA on traits of evolutionary and biomedical importance in a non-human primate model with relevance to humans.

## Introduction

Life history theory seeks to explain how natural selection shapes the allocation of energy among competing physiological demands, typically distributed across growth, reproduction, and somatic maintenance (1-5). Comparative work across the tree of life supports clear trade-offs between these domains: species that grow faster, mature earlier, and reproduce frequently typically live shorter lives, while species that grow slower, mature later, and reproduce less frequently typically live longer lives (6-8). These suites of life history traits are thought to evolve in response to socio-ecological conditions that determine a species’ optimal “strategy” (9, 10). However, how environmental experiences influence suites of life history traits and trade-offs *within* species is less understood (11), but equally important, since this is the intra-specific variation on which natural selection operates (12, 13). Empirical studies demonstrate that early life adversity (ELA; e.g., parental loss, resource constraint, social subordination, etc.) can recalibrate aspects of growth and reproduction in humans and other species (14-17), leading to a large theoretical literature on why certain organisms have evolved to be developmentally plastic and how it might matter for fitness (18-22). However, few studies of natural animal populations have been poised to empirically address early life effects on multiple life history domains simultaneously (23-25), leading to gaps in our understanding of: 1) how ELA influences coordination and trade-offs between life history domains; and 2) whether ELA-induced plasticity in life history traits ultimately matters for fitness.

In humans, recent work has addressed how ELA shapes individual life history traits, but findings remain somewhat inconsistent and challenging to interpret. For example, in humans, adverse early life *social* conditions are generally associated with younger ages at puberty, increased fertility, and higher body weight in adulthood (26-32), while adverse *nutritional* conditions can be associated with shorter stature, low body weight, and either earlier or later reproductive maturation (33-35). Earlier reproduction in humans that experience ELA has sometimes been interpreted as an adaptive life history strategy to maximize reproductive success, in which ELA-exposed individuals prioritize accelerated reproduction over other biological processes in anticipation of early death (36-40). However, this observation, its interpretation, and whether accelerated reproduction following ELA does in fact maximize reproductive success remains debated (41-43). More generally, it remains challenging to understand how much ELA directly influences reproductive biology in humans, because adverse childhood experiences are often confounded by sociocultural factors including access to healthcare and birth control (44, 45), education (46, 47), and retrospective reporting bias of early life experiences (48, 49).

Studies of non-human animals, especially long-lived primates, can circumvent challenges faced by human studies while still providing a biologically relevant model for our species (50). However, identifying ELA effects on reproduction and other life history traits, as well as trade-offs between them, in natural primate populations can be challenging: very few datasets include information on early life environments as well as lifetime growth, reproduction, or other fitness-relevant outcomes. Nevertheless, some progress has been made (51-58). For example, in a long-term study population of baboons, early life drought predicts shorter limb length (59) and cumulative ELA predicts lifetime reproductive success and survival (60-62), yet cumulative ELA is not consistently predictive of the timing or pace of reproduction (43). However, considerably less work has addressed how ELA may induce life history trade-offs with fitness consequences (63); in other words, most existing studies examine either growth or reproduction, leaving a major gap in our understanding of how early life adversity shapes energy allocation between the two.

In this study, we investigate how life history traits are influenced by ELA, whether ELA magnifies trade-offs between life history domains, and how variations in life history across individuals contribute to lifetime reproductive success, a component of evolutionary fitness. To do so, we compare patterns of ELA, growth, and reproduction in a long-term, free-ranging study population of rhesus macaques (*Macaca mulatta*) living on the island of Cayo Santiago off the coast of Puerto Rico. This population of approximately 1,700 individuals is managed by the Caribbean Primate Research Center (CPRC) of the University of Puerto Rico and is regularly monitored by the CPRC and individual researchers who record longitudinal social, demographic, life history, and body condition data. Using a cross-sectional design and established measures of ELA, we investigate 1) how ELA influences growth outcomes (body mass and body size as proxied by trunk and limb lengths), the timing of reproductive maturation, and reproductive rate in a large dataset of 2,100 females monitored between 1960-2021. We also test 2) whether ELA amplifies negative correlations between growth and reproduction and 3) how ELA-impacted growth and reproductive traits relate to lifetime reproductive success. We predict that ELA leads to slower growth and delayed reproduction, based on observations in the study population (57, 64) as well as other wild, long-lived mammals (15, 25). While it is contrary to much of the human literature that suggests ELA facilitates earlier reproductive maturity (26, 27), we suspect that coordinated constraints and intensified trade-offs are generally more common (Figure 1) Together, this work provides a rare test of how early life adversity shapes investment in competing life history domains in a long lived primate.

**Figure 1.**
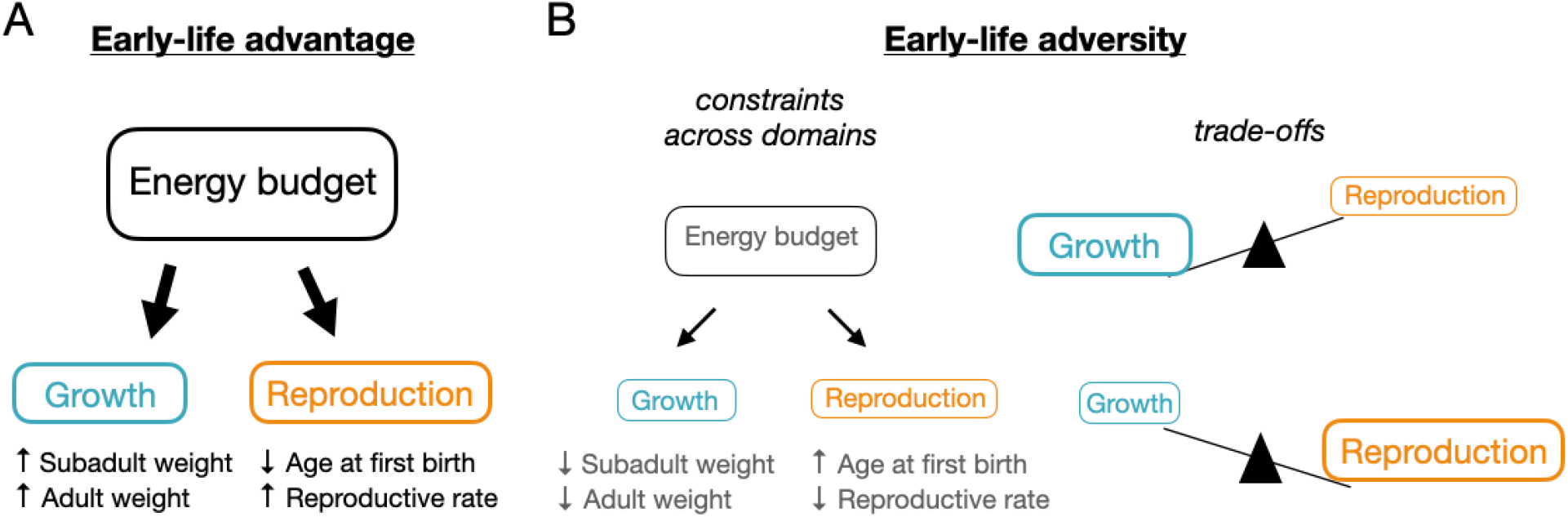
Conceptual models of energetic investment into growth and reproduction. **(A)** Advantageous early life conditions may support high simultaneous investment into both growth and reproduction, leading to greater body weights, an earlier age at first birth, and a faster reproductive rate. **(B)** Adverse early life conditions may be associated with energetic constraints across both growth and reproduction, leading to lower body weights, a later age at first birth, and a slower reproductive rate. Conversely, adversity may be associated with energetic trade-offs between life history domains, wherein enhanced investment into one domain is associated with reduced investment into the other.

## Results

### Early life adversity slows growth and delays reproductive maturation

Using long-term data from the Cayo Santiago rhesus macaque population, we determined age at first birth and reproductive rate for 2,100 females from 1960-2021 (Figure S1-2). Macaques are seasonal breeders; females typically have their first birth at 4-5 years old and can (but do not always) give birth to an offspring each year (65). As such, reproductive rate can be calculated as the proportion of years in which a female gave birth following their first birth season and their total surviving offspring calculated as the number that reached reproductive maturity (4 years old). From these 2,100 females with reproductive histories, we also obtained body weights collected during annual trapping efforts for a subset of 861 females (n=1,339 measurements; Figure S3), and limb/trunk length measurements from 325 females (n=460 measurements). Lastly, we calculated seven established sources of ELA—early maternal loss, maternal primiparity, having a close in age younger sibling, small kin network size, large group size, low maternal dominance, and experiencing a natural disaster—as well as cumulative adversity (the number of experienced individual adversities; Figure S4; see Table S1 for detailed ELA descriptions). These variables have been shown to predict survival in this and other non-human primate populations (61, 66). Notably, we found that our seven individual forms of ELA were not correlated (Figure S5), providing a unique opportunity to isolate the effects of different types of ELA, which is generally not feasible in human studies where adversities often co-occur (67). When running models across multiple ELAs, we use a false discovery rate (FDR) correction (68); we report significant associations as those with an FDR < 0.1 and nominally significant associations as those with an unadjusted p < 0.05.

First, we aimed to understand whether ELA influences life history traits, focusing on measures of growth and reproductive timing during development. We modeled the effect of ELA on 1) sub-adult body weight (< 4 years old) and 2) the age at which a female gave birth to her first surviving offspring. We found that two sources of ELA modified the relationship between age and body weight during early life (Figure 2A; Table S2): females with a competing younger sibling had lower sub-adult body mass compared to females without a competing sibling (n=612, estimate= -0.17, standard error (SE)= 0.04, FDR=2.2x10^-4^), and females born into large groups, a potential form of adversity due to enhanced intra-group competition, were heavier as sub-adults compared to females born into smaller groups (Figure 2B; n=612, estimate=0.08, SE=0.02, FDR=2.5x10^-4^).

**Figure 2.**
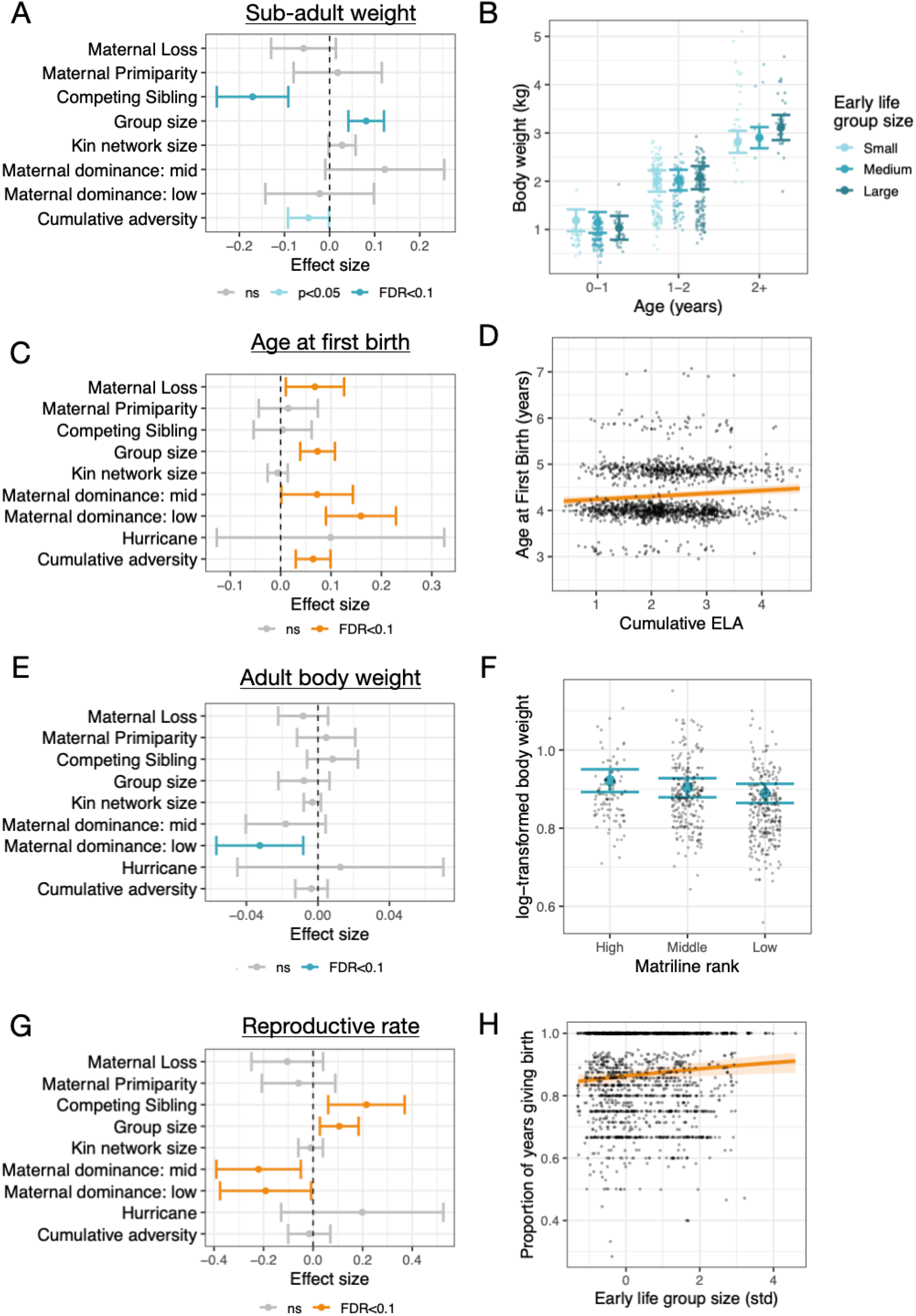
Early life adversity predicts body size and reproduction across the lifecourse. **(A)** Model results depicting the effect of each adversity on the interaction between age and ELA in predicting sub-adult body weight. **(B)** Effect of early life group size on body weight across development. **(C)** Model results depicting the effect of each adversity in predicting a female’s age at first birth. **(D)** Greater cumulative ELA is associated with a later age at first birth. **(E)** Model results depicting the effect of each adversity on adult body weight. **(F)** Lower adult body weight is associated with being born into a larger social group. **(G)** Model results depicting the effect of each adversity on reproductive rate. **(H)** Faster reproductive rate is associated with being born into a larger social group. A, C, E, F: Points represent model estimates, error bars represent the standard error of the estimate, and color represents the significance threshold. B, D, F, G: Large points/colored lines represent model estimates, error bars/shading represent the 95% confidence interval, and small points represent raw data.

Second, we found that three different forms of ELA were associated with delayed reproductive maturation (Figure 2C; Table S2). Females who experienced early maternal loss (n= 2100, estimate= 0.07, SE=0.03, FDR=0.048), were born into larger groups (n= 2098; estimate=0.07, SE=0.02, FDR=1.7 x 10^-4^), or were born to mothers in medium or low ranking matrilines (n= 1649; medium rank v high rank estimate=0.07, SE=0.04, FDR=0.083; low rank v high rank estimate=0.16, SE=0.04, FDR=7.1x10^-5^), started reproducing at older ages. Specifically, individuals who experienced maternal loss, were in the top 90th percentile of group sizes, or were in low ranking matrilines were predicted to start reproduction 0.8, 2.4, and 1.9 months later than their counterparts that did not experience maternal loss, were in the bottom 10th percentile of group sizes, or were in high ranking matrilines, respectively. These results were recapitulated using our cumulative ELA metric, where those experiencing four adversities were predicted to begin reproducing three months later than those experiencing no adversities (Figure 2D; n= 1640; estimate= 0.06, SE= 0.02, FDR=8.0x10^-4^).

### Early life adversity is linked to lower adult body weight and modified reproductive rate

To understand how ELA impacts life history during adulthood, we next tested whether ELA is associated with individual differences in: 1) adult body weight (> 4 years old); 2) adult limb and trunk lengths; and 3) reproductive rate (proportion of birth seasons in which a female gave birth during her reproductive career). We found that one form of ELA was associated with adult body weight (Figure 2E; Table S2): adult body weight was negatively associated with matriline rank (Figure 2F; n=605, estimate=-0.03, SE=0.01, FDR=0.08), such that individuals born into lower ranking matrilines were predicted to weigh 0.6kg (1.3 pounds) lighter in adulthood than those born into high ranking matrilines. In contrast, adult arm, leg, and trunk lengths were not associated with any form of ELA (Table S2).

Three different forms of ELA were associated with reproductive rate (Figure 2G; Table S2): females born into mid and low ranking matrilines had a lower probability of giving birth each year (i.e., slower reproductive rate) compared to females born into high ranking matrilines (n=1291; medium rank v high rank estimate= -0.22, SE=0.09, FDR= 0.034; low rank v high rank estimate=-0.19, SE=0.09, FDR=0.09), equating to a 2.2 percentage point lower probability in low versus high ranking females. Conversely, females who had a competing younger sibling (Figure 2H; n=1419; estimate=0.22, SE=0.08, FDR=0.034), or who were born into larger groups (n=1417; estimate=0.1, SE=0.04, FDR=0.034) had a higher probability of giving birth each year (i.e., faster reproductive rate).

### Early life adversity enhances trade-offs between growth and reproduction

To assess whether ELA may alter an individual’s relative investment into growth and reproduction, we first confirmed that these processes are linked during development. We found that sub-adult body size negatively predicted age at first birth, such that females who were small for their age experienced their first birth later than their larger peers (Figure 3A; Table S3; n=201, estimate= -0.42, SE= 0.14, p=0.007). Next, we tested whether ELA modified this relationship. Though no interaction terms reached statistical significance following multiple hypothesis testing correction, adversity in the form of large early life group size trended toward strengthening the negative association between age at first birth and sub-adult body weight. Specifically, females born into large groups who were small for their age trended toward delayed reproduction even more than similarly sized females from smaller groups (Figure 3B; Table S3; n=201, estimate=-0.3, SE= 0.14, p=0.04, FDR=0.31).

**Figure 3.**
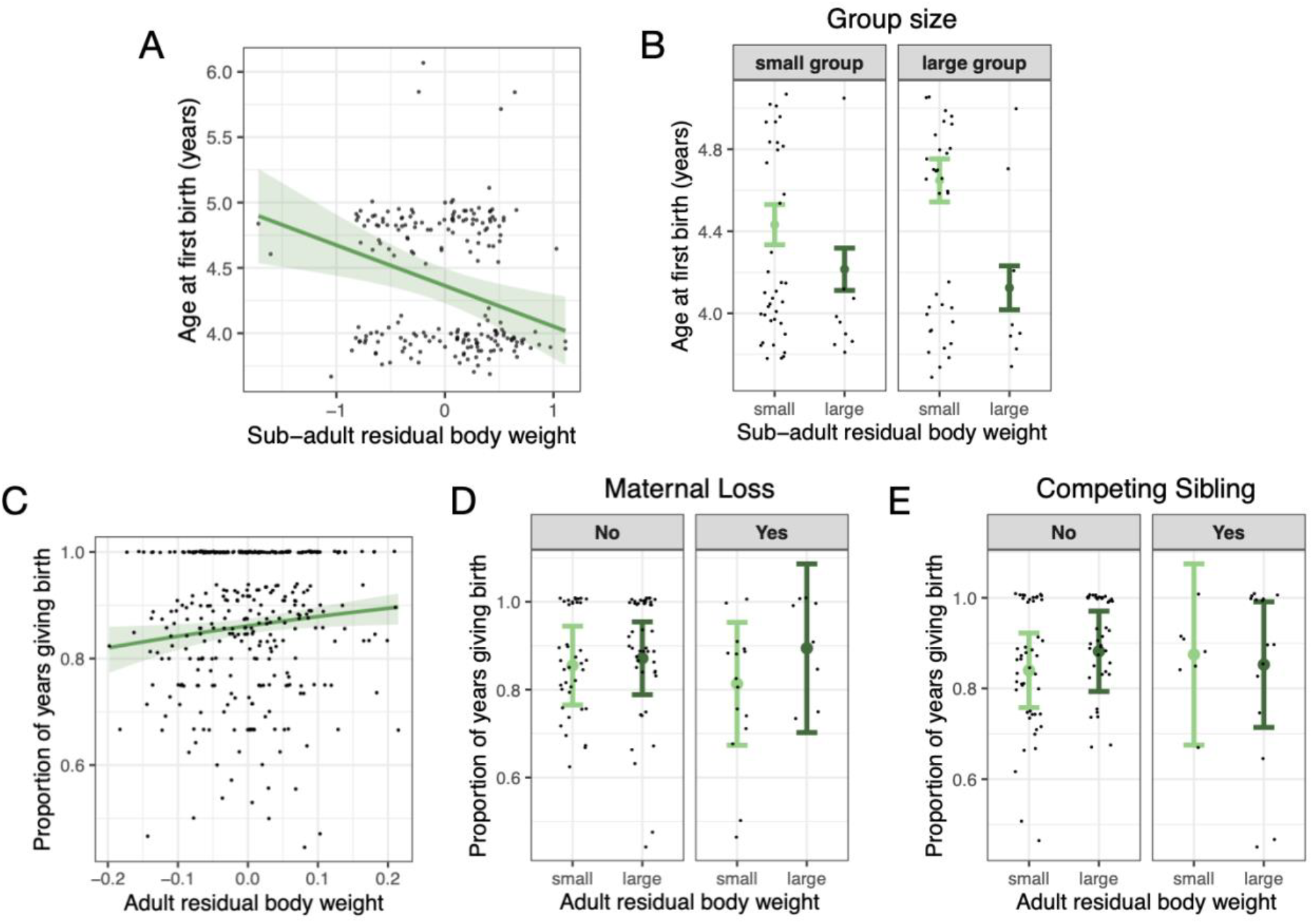
Early life adversity exacerbates and modifies trade-offs between growth and reproduction. **(A)** Negative relationship between sub-adult body weight and age at first birth, indicating that larger individuals are predicted to start reproducing earlier. **(B)** Being born into a large group exacerbates the negative relationship between sub-adult body weight and age at first birth. **(C)** Positive relationship between adult body weight and the proportion of years a female gives birth, indicating that larger individuals are predicted to have a faster reproductive rate. **(D)** Experiencing maternal loss exacerbates the negative relationship between body weight and reproductive rate, and **(E)** having a competing younger sibling *modifies* the relationship between body weight and reproductive rate, such that small females who experienced adversity reproduce faster than similar sized females that did not experience adversity. A & C: Green line represents the model prediction, shading represents the standard error, and points represent the raw data. B, D & E: Colored points and error bars represent the model prediction and standard error at +/- 1SD of mean body weight (small vs large) and +/- 1SD of mean group size (small group vs large group). Black points represent raw data.

We also examined whether ELA might alter the relative investment into growth and reproduction in adulthood by testing whether adult body weight predicts reproductive rate. We found a positive relationship, such that females who were larger than average had faster reproductive rates (Figure 3C; Table S3; n=353, estimate=1.55, SE= 0.68, p=0.02). Next, we tested whether ELA modifies this relationship. Similar to the pattern observed during development, we found that two forms of ELA, maternal loss and low matriline rank, exacerbated the trade-off between body size and reproductive rate: females who experienced ELA who were small for their age gave birth less frequently than similarly sized females who did not experience ELA (Figure 3D; Table S3; maternal loss: n=353, estimate= 3.42, SE=1.69, FDR=0.09; matriline rank: n=299, estimate=3.9, SE=1.9, FDR=0.09). Surprisingly, we found that two different forms of ELA, having a competing sibling and being born to a primiparous mother, *reduced* the trade-off between body size and reproductive rate: females who experienced these adversities that were small for their age gave birth more frequently than similar sized females who did not experience these adversities (Figure 3E; Table S3; competing sibling: n=353, estimate= -3.55, SE=1.67, FDR=0.09; primiparity: n=353, estimate=-5.68, SE=2.0, FDR=0.03).

### Variation in life history predicts lifetime reproductive success

Lastly, we aimed to understand whether ELA influenced life history traits that matter for a component of evolutionary fitness: lifetime reproductive success. To do so, we first tested whether a female’s age-adjusted body weight (including both adult and sub-adult weights), age at first birth, or reproductive rate predicted their number of offspring that survived to reproductive maturity (4 years old), focusing on females for which we have data from their entire reproductive career (i.e., who were born and died naturally on the island). We found no evidence for a relationship between age-adjusted body weight and number of surviving offspring. However, we did find that females who started reproducing earlier and who reproduced at a moderate rate had a greater number of offspring who survived to reproductive maturity (Figure 4B&C; Table S4; age first birth: n=279, estimate=-0.66, SE=0.27, FDR=0.03; reproductive rate: n=279, estimate=-5.51, SE=1.28, FDR=1.2x10^-7^).

**Figure 4.**
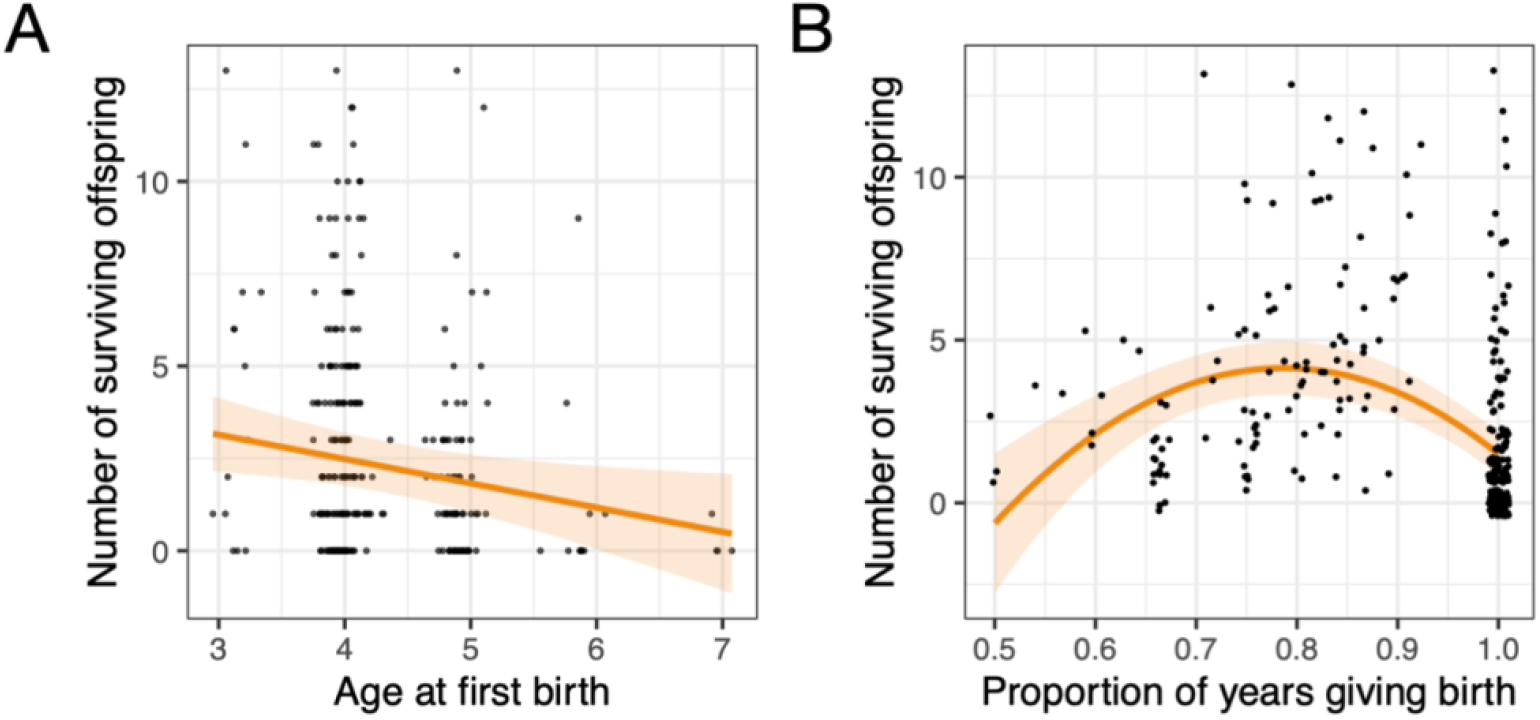
Variation in reproductive life history impacts a proxy of fitness. **(A)** younger age at first birth and **(B)** a moderate reproductive rate are associated with a greater number of surviving offspring. Orange lines represent the model prediction, shading represents the standard error, and points represent the raw data. These analyses were restricted to females who completed their full reproductive careers and therefore represent a subset of the overall dataset.

## Discussion

Here, we aimed to understand trade-offs between growth and reproduction following ELA in the free-ranging rhesus macaques of Cayo Santiago. Exposure to ELA was linked to both body mass and reproductive effort, generally resulting in smaller females who reached reproductive maturity later and reproduced either faster or slower than non-ELA exposed females. Data on body weight and reproduction of the same female allowed us to test for trade-offs between these life history domains: we observed that females who were small for their age started reproducing later and maintained slower reproductive rates in adulthood, as observed in other non-human primates (69). While some adversities exacerbated this trade-off, other adversities decoupled the relationship, such that adversity-exposed females who were small were reproducing *faster* than those who were large. Lastly, we found that variations in the timing of reproductive maturity and reproductive rate contributed to a female’s reproductive success, suggesting that ELA-induced plasticity in life history might matter for fitness.

Our findings provide compelling evidence that ELA is associated with a delayed reproductive debut in female rhesus macaques. This result offers valuable insight into the divided evidence supporting a link between ELA and both an earlier and later age of puberty in humans. In humans, earlier pubertal timing is most consistently observed following experiences of abuse, particularly sexual abuse (70-72), which has been attributed to the broad social stressors that likely co-occur in such environments (73). In contrast, the broad range of early life adversities tested here were linked to a later age at first birth, with none associated with an earlier age at first birth. Given this, we suggest that the link between ELA and early maturation in humans may be rooted in particular adversity types (i.e., sexual or psychological abuse) that are not broadly experienced in other non-human primate species. As such, physiological acceleration of reproductive timing following early life sexual trauma in humans may not reflect an evolutionarily conserved adaptation, but a response to uniquely human social environments and contexts.

Contrary to our predictions, our results also suggest that some adversities (e.g., large early life group size and having a competing sibling) are associated with an accelerated reproductive rate. As seasonal breeders, females whose offspring die soon after birth resume cycling faster and are more likely to conceive during the following mating season than females whose offspring survive (74). Accordingly, we observe that females with the highest reproductive rates have fewer surviving offspring than females with moderate reproductive rates. In this context, ELA may predict faster reproduction due to lower offspring survival, underscoring the necessity of evaluating the life history consequences of ELA in tandem with measures of fitness. Furthermore, these results suggest a potential trade-off between offspring quality and quantity, a phenomenon that is well-documented within and across mammalian species (75, 76), though limited previous work has identified impacts of ELA (77). In humans, childhood adversity is associated with reduced gestational investment and short inter-pregnancy intervals (29, 78), both of which are linked to higher offspring mortality (79-81). Our results provide corroborating evidence of a quantity over quality reproductive strategy following ELA in a long-lived non-human primate species with similar reproductive patterns to humans, including singleton births and interbirth intervals that predict offspring survival.

Importantly, our matched body weight and reproductive data types have enabled us to identify within-individual trade-offs between growth and reproduction, which appear to operate differently depending on the specific type of adversity experienced. For example, small-bodied females who lost their mothers reproduced more slowly than equally small females who did not experience maternal loss, suggesting that this form of ELA can impose correlated constraints across both life history domains. In contrast, other ELAs appear to modify this trade-off in the opposite direction, such that ELA-exposed females of small body size exhibited *increased* reproductive output. Across mammals, high reproductive rates are often linked with smaller body size, but within a species this relationship is typically reversed, with larger females experiencing greater reproductive output and success (82, 83). In the Cayo Santiago macaques, however, maximal reproductive rates were associated with diminished reproductive success, likely due to the link between offspring loss and a rapid return to fertility. Thus, while different ELAs may influence the growth–reproduction trade-off in opposing ways, both likely result in suboptimal reproductive success. This underscores the idea that ELA generally imposes constraints on optimal life history trajectories which may not be evident when investigating reproduction or growth in isolation or without measures of fitness.

Despite its contributions, this study has several important limitations. First, although we use body size measurements to approximate growth, our data are cross-sectional. Future longitudinal studies will be valuable for understanding within-individual trade-offs between growth and reproduction across the life course. Second, while we were able to statistically control for major aspects of the current social environment, such as a female’s current group size and kin network size, this type of adjustment was not feasible for all forms of adversity. For example, matriline rank is socially inherited from mothers to daughters and relatively stable over time (84), meaning early life matriline rank is often correlated with later-life rank. Third, the Cayo Santiago rhesus macaques are partly provisioned (85), a factor that inherently alters the resource landscape on the island. As a result, ELAs which are presumed to contribute to differential resource access may be dampened in this population in comparison to fully wild populations. For instance, whereas a previous study in baboons found that early life drought was associated with reduced limb length (59), we did not observe a comparable relationship in the Cayo population. Finally, this work does not directly examine the most commonly presumed third life history domain, somatic maintenance, and would benefit, for example, from measures of immune investment. Studies incorporating additional molecular measures to clarify how this third domain of life history integrates and experiences trade-offs with growth and reproduction would be extremely fruitful (86, 87).

These findings offer new insight into how adversity shapes energy allocation between life history domains by characterizing within-individual trade-offs between growth and reproduction in conjunction with fitness outcomes in a long-lived mammal. While our study focuses on females, ELA is often predicted to have even stronger effects on males, underscoring the need for future research on sex-specific responses (88-90). Unlike studies of humans where different adversities often co-occur and are confounded by systemic social inequities, adversities in our population were largely uncorrelated, allowing us to identify unique responses to distinct adversity types. Accordingly, we did not find strong effects of cumulative adversity across life history processes, as has previously been shown in studies of survival (61, 66, 91), suggesting that different ELAs may have pathway-specific rather than additive effects. Future molecular studies will be poised to answer important questions about whether different adversities converge on shared biological mechanisms or act through distinct tissues or systems (e.g., the HPA axis vs. nutritional pathways) (92). Finally, our results challenge the idea that earlier menarche following adversity in humans reflects a conserved primate strategy, and instead may be a unique response to trauma that cannot be meaningfully approximated in naturalistic non-human primate populations. Instead, our results support a model in which adversity constrains both growth and reproduction, pushing individuals away from fitness-optimal life histories.

## Materials and methods

### Study population

This study leverages long-term data from the free-ranging population of rhesus macaques on Cayo Santiago, a 5.2 hectare island located 1 km off the Southeast coast of Puerto Rico. This colony originated from 409 rhesus macaques of Indian origin introduced to the island in 1938 and systematically observed since the 1960s (93). The population has fluctuated in size across decades (94) and now comprises approximately 1,700 individuals. This population exhibits reproductive seasonality with an approximate 1 year interval, with many females giving birth to a single offspring each year (95, 96). The Cayo Santiago macaques have no natural predators on the island and thus we do not assess predation risk (an ecological factor known to impact life history in some species), but instead focus on aspects of the maternal and group environment as well as exposure to extreme weather such as major hurricanes which are known to hit Puerto Rico at Category 4 and 5 intensity. They are provided with fresh drinking water and are partially provisioned with monkey chow, although they still forage on the natural vegetation on the island (85). Although this population is provisioned, they exhibit species-typical behaviors, including the establishment, maintenance and natural fissioning of social groups, dominance hierarchies, as well as similar somatic development in terms of lifespan and body size compared to their wild counterparts (97, 98). Each year, the CPRC traps and releases yearlings, marking them with unique tattoos to facilitate individual recognition of all individuals on the island. Additionally, the CPRC periodically removes subsets of individuals from the island to mitigate overpopulation. All data used in this study was collected using protocols approved by the University of Puerto Rico Institutional Animal Care and Use Committee (protocol # A400117)

### Demographic and reproduction data

Since 1956, the CPRC records all changes in island demography including births, deaths, migration and group composition on a nearly daily basis. We limited our analyses to individuals born in 1960 and after because no systematic records are available prior to 1960. We removed outliers, consisting of instances in which the age of first birth or average IBI was 5 standard deviations above or below the mean, suggesting an error in measurement. After these filtering steps, our final dataset included 2,100 individual mothers. In this dataset, mean age at first birth was 4.35 years old (range 2.9-7.1 years), and the average female gave birth 7.4 times (range 1-20). Although individuals are monitored nearly daily, early reproductive failures—such as abortions or very early infant deaths—may go undetected, so recorded births per female may be slightly underestimated. To calculate the number of surviving offspring for each female, we limited our dataset to females for which we have data on their entire reproductive career (i.e., who were born and died naturally on the island and were not removed during CPRC population management efforts: (96)) and whose offspring all either naturally died on the island, were removed from the island but after the age of 4 years old, or were still living. This left us with a subset of 279 females for which we had data for their entire reproductive career, with females averaging 2.7 offspring surviving to reproductive age (range: 0-13).

### Growth data

Body weights and body size measurements (i.e., trunk and limb lengths) from adult individuals (> 4 years old) were taken from 2019-2024 during the annual trap and release period on Cayo Santiago as part of a study on the longitudinal effects of aging. Body weights of sub-adult individuals (< 4 years old) were collected from 2005-2011 as part of projects led by author A.W. which have been previously published (99, 100). Additionally, body weight and body size measurements were obtained from individuals removed from the population following CPRC population management efforts in 2016, 2018, 2019, and 2021. In total, our growth dataset consisted of 1,339 body weights from 861 unique individuals (mean=1.6 per female, range=1-7) and 460 body size measurements taken from 325 unique individuals (mean=1.4 per female, range 1-6). Although some individuals with sub-adult body weights were also measured in adulthood (n=84), the majority were not. See Supplementary Methods for more details on body size measurements.

### Early life adversity data

We quantified seven sources of ELA: early maternal loss, maternal primiparity, having a close in age sibling, small maternal kin network size, large group size, low matrilineal dominance rank, and experiencing a natural disaster (i.e., a major hurricane). These sources of ELA have been well-established in other non-human primate models and have been linked to survival in this population (43, 61, 66, 101). These forms of adversity can impact an infant’s access to adequate social and/or nutritional support and are homologous with sources of adversity identified in humans (102). Following Tung et al. (61), we used these metrics to calculate cumulative adversity exposure as the sum of all individual adversities an individual experienced. See Supplementary Methods for more details.

### Current environment data

To better isolate the effects of ELA, we calculated two aspects of a female’s current environment to use as covariates in statistical models of adult reproduction and growth: group size and kin network size. Current group size consisted of the number of adult males and females in an individual’s social group on either 1) the date of body weight measurement, or 2) on the date that a female gave birth averaged over all births, for adult body weight and reproductive rate models, respectively. Likewise, we calculated current kin network sizes as the number of close adult female relatives (r ≥ 0.063) in the group on the date of body weight measurement or the average number of close adult female relatives averaged across each of the days in which a female gave birth.

### Statistical methods

To understand how ELA influences age at first birth, subadult body weight, and adult body weight, limb length, and trunk length, we fit linear mixed-effects models (LMMs) including each ELA as a predictor in a separate model and natal group and birth year as random effects. To assess the relationship between ELA and reproductive rate, we fit generalized linear mixed effects models (GLMMs) with a binomial error distribution and logit link. The response was the number of births relative to the number of birth seasons a female was alive for during her reproductive career. Each model included one ELA predictor, standardized adult group size and kin network size as covariates, and birth year and natal group as random effects. To model the relationship between ELA and growth, as proxied by subadult body weight, we tested whether ELA modified the relationship between age and subadult body weight (modeled as an interaction effect). Due to the timing of sampling, there were no individuals for which we had early life body weight that experienced a hurricane in their first year of life, and thus we did not assess whether hurricane exposure modified the relationship between age and subadult body weight. To model the relationship between ELA and adult body weight and size, we used a log-log model in which both the response variable and the covariate age were log transformed, to account for the nonlinear relationship between body weight and age in adulthood and following previous modeling studies of adult body size in baboons (59). All body weight and trunk length models included age as a quadratic term which provided a significantly better fit than those with age modeled as a linear covariate, as indicated by lower AIC values. We included standardized adult group size and kin network size as covariates and animal ID as a random effect. We did not include current environment variables in the age at first birth or subadult body weight models as females typically remain in their natal groups and so their social environment at their age of first birth will still strongly reflect the birth environment. We adjusted all p-values for models predicting a given life history metric using the Benjamini-Hochberg false discovery rate (FDR) correction (68). We report significant associations as those with an FDR < 0.1, a common cutoff in ecological studies indicative of a 10% false positive rate across tests (103).

To investigate trade-offs between growth and reproduction, we fit LMMs testing for the effect of subadult residual body weight (weight controlled for age, as described above) on age at first birth, and adult residual body weight on the proportion of birth seasons in which a female gave birth. Here, we report unadjusted p-values, as each test is performed only once. We then tested for an interaction effect between residual body weight (subadult or adult) and ELA in predicting each reproductive metric, and controlled for FDR (68). The majority of individuals were weighed only once, and thus we did not include individual ID as a random effect. Instead, we included only one randomly chosen body weight measurement per individual, resulting in a total of 201 individuals with subadult body weight and age at first birth and 353 individuals with adult body weight and reproductive rate data.

To understand how variation in life history may contribute to evolutionary fitness in this population, we fit LMMs including one life history metric as the predictor variable, number of offspring surviving to 4 years old as the response variable, and birth season and natal group as random effects. To test for non-linear effects, we ran each model with the life history metric as a linear term and one including both a linear and quadratic term and used AIC to test for model fit. We included in this analysis only females who died naturally on the island (i.e., were not removed by colony management) so that the number of surviving offspring represents a female’s entire reproductive career. Because the majority of body weight measurements were taken within the last 5 years many of these females are still alive, and thus only 52 individuals with body weight data were deceased and therefore had information on their full reproductive career. Due to this small sample size, we combined both subadult and adult body weights into one analysis. To account for differences in weight due to age at sampling, we used age-adjusted weights to capture whether individuals were heavier or lighter than expected for their age. An individual’s age-adjusted weight was their residual value extracted from a linear model wherein age predicted body weight. Due to linear growth patterns in sub adulthood, we used raw age as the predictor variable and body weight as the outcome variable for individuals less than 4 years old. To account for the non-linear relationship between body weight and age in adulthood, we used a quadratic log-log model (log-transformed response and quadratic log-transformed predictor) following Levy et al. (59) to model the effects of age on body weight and calculate age-adjusted weight for individuals over 4 years old.

## Data Availability

All data reported in this paper are publicly available on Zenodo (https://doi.org/10.5281/zenodo.17095402), and all code is available on RMP’s personal github (https://github.com/rachpetersen/Cayo_ELA_LifeHistory)

## Author Contributions

R.M.P., S.K.P., J.P.H., and A.J.L. designed research, A.W., C.M.T., S.K.P., L.J.N.B., S.A.W., S.C.A., A.N.R., S.E.B.S., C.B.R.U., and A.V.R.L. contributed data, R.M.P., S.K.P., J.P.H., and A.J.L. analyzed data, R.M.P., S.K.P., J.P.H., and A.J.L. wrote the paper. All authors read and edited the paper.

## Competing Interest Statement

The authors declare no competing interests

## Classification

Biological Sciences: Evolution

## Funding sources

National Science Foundation SBE Postdoctoral Research Fellowship 2313953 (RMP), NSF-SMA-2105307 (SKP), R21-AG078554 (AJL), R00-AG051764 (NSM), R01-AG060931 (NSM), R01-MH096875 (CBRU), R01-MH089484 (CBRU), R01-MH118203 (CBRU), R01-AG084706 (JPH), R56-AG071023 (JPH), NSF-RAPID-1800558 (JPH/SCA), German Research Foundation (DFG) Emmy Noether group (grant numbers WI 1808/1-1, 1-2, 2-1, 3-1 to AW); The CPRC is supported by the Office of Research Infrastructure Programs (ORIP) of the National Institute of Health (NIH) through grant number P40 OD012217.

## Acknowledgements

We would like to thank members of the Caribbean Primate Research Center at the University of Puerto Rico for the long-term collection and curation of the data presented here.

## Notes

### Competing Interest Statement

The authors have declared no competing interest.

### Summary of Updates

This version of the manuscript has been revised to update the author list.

## References

1. A. J. Van Noordwijk, G. De Jong, Acquisition and allocation of resources: their influence on variation in life history tactics. The American Naturalist 128, 137–142 (1986).

2. E. L. Charnov, Evolution of life history variation among female mammals. Proceedings of the National Academy of Sciences 88, 1134–1137 (1991).

3. S. C. Stearns, The evolution of life histories (Oxford university press, 1998).

4. J. Kozłowski, Optimal allocation of resources to growth and reproduction: implications for age and size at maturity. Trends in ecology & evolution 7, 15–19 (1992).

5. J. R. Burger, C. Hou, J. H. Brown, Toward a metabolic theory of life history. Proceedings of the National Academy of Sciences 116, 26653–26661 (2019).

6. E. R. Pianka, On r-and K-selection. The american naturalist 104, 592–597 (1970).

7. D. E. Promislow, P. H. Harvey, Living fast and dying young: A comparative analysis of life-history variation among mammals. Journal of Zoology 220, 417–437 (1990).

8. S. Hamel et al., Fitness costs of reproduction depend on life speed: empirical evidence from mammalian populations. Ecology letters 13, 915–935 (2010).

9. L. Marty, U. Dieckmann, M.-J. Rochet, B. Ernande, Impact of environmental covariation in growth and mortality on evolving maturation reaction norms. The American Naturalist 177, E98–E118 (2011).

10. T. J. Kawecki, S. C. Stearns, The evolution of life histories in spatially heterogeneous environments: optimal reaction norms revisited. Evolutionary Ecology 7, 155–174 (1993).

11. M. Galipaud, H. Kokko, Adaptation and plasticity in life-history theory: How to derive predictions. Evolution and Human Behavior 41, 493–501 (2020).

12. S. C. Antón, C. W. Kuzawa, Early Homo, plasticity and the extended evolutionary synthesis. Interface Focus 7, 20170004 (2017).

13. M. J. West-Eberhard, Developmental plasticity and evolution (Oxford University Press, 2003).

14. D. Meikle, M. Westberg, Maternal nutrition and reproduction of daughters in wild house mice (Mus musculus). REPRODUCTION-CAMBRIDGE- 122, 437–442 (2001).

15. M. Gicquel, M. L. East, H. Hofer, S. Benhaiem, Early-life adversity predicts performance and fitness in a wild social carnivore. Journal of Animal Ecology 91, 2074–2086 (2022).

16. C. W. Kuzawa, Developmental origins of life history: growth, productivity, and reproduction. American Journal of Human Biology 19, 654–661 (2007).

17. S. Descamps, S. Boutin, D. Berteaux, A. G. McAdam, J.-M. Gaillard, Cohort effects in red squirrels: the influence of density, food abundance and temperature on future survival and reproductive success. Journal of Animal Ecology, 305–314 (2008).

18. D. Nettle, M. Bateson, Adaptive developmental plasticity: what is it, how can we recognize it and when can it evolve? Proceedings of the Royal Society B: Biological Sciences 282, 20151005 (2015).

19. D. J. Marshall, T. Uller, When is a maternal effect adaptive? Oikos 116, 1957–1963 (2007).

20. P. D. Gluckman, M. A. Hanson, H. G. Spencer, Predictive adaptive responses and human evolution. Trends in ecology & evolution 20, 527–533 (2005).

21. T. W. Fawcett, W. E. Frankenhuis, Adaptive explanations for sensitive windows in development. Frontiers in Zoology 12, S3 (2015).

22. B. Fischer, G. S. van Doorn, U. Dieckmann, B. Taborsky, The evolution of age-dependent plasticity. The American Naturalist 183, 108–125 (2014).

23. M. C. Forchhammer, T. H. Clutton-Brock, J. Lindström, S. D. Albon, Climate and population density induce long-term cohort variation in a northern ungulate. Journal of Animal Ecology 70, 721–729 (2001).

24. S. Albon, T. Clutton-Brock, F. Guinness, Early development and population dynamics in red deer. II. Density-independent effects and cohort variation. The Journal of Animal Ecology, 69–81 (1987).

25. G. Pigeon, F. Pelletier, Direct and indirect effects of early-life environment on lifetime fitness of bighorn ewes. Proceedings of the Royal Society B: Biological Sciences 285, 20171935 (2018).

26. A. K. Pesonen et al., Reproductive traits following a parent–child separation trauma during childhood: a natural experiment during World War II. American Journal of Human Biology: The Official Journal of the Human Biology Association 20, 345–351 (2008).

27. M. Klimek et al., Early-life adversities and later-life reproductive patterns in women with fully traced reproductive history. Scientific Reports 13, 9328 (2023).

28. J. Liimatta et al., Accelerated Early Childhood Growth Is Associated With the Development of Earlier Adrenarche and Puberty. Journal of the Endocrine Society 8, bvae026 (2024).

29. E. A. Holdsworth, A. A. Appleton, Adverse childhood experiences and reproductive strategies in a contemporary US population. American Journal of Physical Anthropology 171, 37–49 (2020).

30. D. Amir, M. R. Jordan, R. G. Bribiescas, A longitudinal assessment of associations between adolescent environment, adversity perception, and economic status on fertility and age of menarche. PLoS One 11, e0155883 (2016).

31. T. Fleischer et al., The relation between childhood adversity and adult obesity in a population-based study in women and men. Scientific reports 11, 14068 (2021).

32. L. K. Elsenburg, K. J. van Wijk, A. C. Liefbroer, N. Smidt, Accumulation of adverse childhood events and overweight in children: a systematic review and meta-analysis. Obesity 25, 820–832 (2017).

33. L. Ibáñez, A. Ferrer, M. V. Marcos, F. R. Hierro, F. de Zegher, Early puberty: rapid progression and reduced final height in girls with low birth weight. Pediatrics 106, e72–e72 (2000).

34. R. Verkauskiene, I. Petraitiene, K. Albertsson Wikland, Puberty in children born small for gestational age. Hormone research in paediatrics 80, 69–77 (2013).

35. J. Yuan, Y. Yu, D. Liu, Y. Sun, Associations between distinct dimensions of early life adversity and accelerated reproductive strategy among middle-aged women in China. American Journal of Obstetrics and Gynecology 226, 104. e101-104. e114 (2022).

36. J. Belsky, L. Steinberg, P. Draper, Childhood experience, interpersonal development, and reproductive strategy: An evolutionary theory of socialization. Child development 62, 647–670 (1991).

37. J. Belsky, The development of human reproductive strategies: Progress and prospects. Current Directions in Psychological Science 21, 310–316 (2012).

38. D. Nettle, W. E. Frankenhuis, I. J. Rickard, The evolution of predictive adaptive responses in human life history. Proceedings of the Royal Society B: Biological Sciences 280, 20131343 (2013).

39. J. S. Chisholm, J. A. Quinlivan, R. W. Petersen, D. A. Coall, Early stress predicts age at menarche and first birth, adult attachment, and expected lifespan. Human nature 16, 233–265 (2005).

40. R. S. Walker, M. J. Hamilton, Life-history consequences of density dependence and the evolution of human body size. Current Anthropology 49, 115–122 (2008).

41. B. S. Low, A. Hazel, N. Parker, K. B. Welch, Influences on women’s reproductive lives: Unexpected ecological underpinnings. Cross-Cultural Research 42, 201–219 (2008).

42. B. J. Ellis, A. J. Figueredo, B. H. Brumbach, G. L. Schlomer, Fundamental dimensions of environmental risk: The impact of harsh versus unpredictable environments on the evolution and development of life history strategies. Human nature 20, 204–268 (2009).

43. C. J. Weibel, J. Tung, S. C. Alberts, E. A. Archie, Accelerated reproduction is not an adaptive response to early-life adversity in wild baboons. Proceedings of the National Academy of Sciences 117, 24909–24919 (2020).

44. H.E. Alcalá, A. Valdez-Dadia, O. S. von Ehrenstein, Adverse childhood experiences and access and utilization of health care. Journal of Public Health 40, 684–692 (2018).

45. A. Srivastav, C. L. Richard, C. Kipp, M. Strompolis, K. White, Racial/ethnic disparities in health care access are associated with adverse childhood experiences. Journal of Racial and Ethnic Health Disparities 7, 1225–1233 (2020).

46. E. Crouch, E. Radcliff, P. Hung, K. Bennett, Challenges to school success and the role of adverse childhood experiences. Academic pediatrics 19, 899–907 (2019).

47. L. C. Houtepen et al., Associations of adverse childhood experiences with educational attainment and adolescent health and the role of family and socioeconomic factors: a prospective cohort study in the UK. PLoS medicine 17, e1003031 (2020).

48. A. Reuben et al., Lest we forget: comparing retrospective and prospective assessments of adverse childhood experiences in the prediction of adult health. Journal of Child Psychology and Psychiatry 57, 1103–1112 (2016).

49. S. N. Naicker, S. A. Norris, L. M. Richter, Secondary analysis of retrospective and prospective reports of adverse childhood experiences and mental health in young adulthood: Filtered through recent stressors. EClinicalMedicine 40 (2021).

50. K. A. Phillips et al., Why primate models matter. American journal of primatology 76, 801–827 (2014).

51. F. Pittet, C. Johnson, K. Hinde, Age at reproductive debut: Developmental predictors and consequences for lactation, infant mass, and subsequent reproduction in rhesus macaques (Macaca mulatta). American Journal of Physical Anthropology 164, 457–476 (2017).

52. S. K. Patterson et al., Effects of early life adversity on maternal effort and glucocorticoids in wild olive baboons. Behavioral Ecology & Sociobiology 75, 1–18 (2021).

53. J. Altmann, S. C. Alberts, Growth rates in a wild primate population: ecological influences and maternal effects. Behavioral Ecology & Sociobiology 57, 490–501 (2005).

54. E. Neves et al., Influence of environmental conditions on population growth and age-specific vital rates of a long-lived primate species in two contrasted habitats. Population Ecology 66, 71–85 (2024).

55. A. Berghänel, M. Heistermann, O. Schülke, J. Ostner, Prenatal stress effects in a wild, long-lived primate: predictive adaptive responses in an unpredictable environment. Proceedings of the Royal Society B: Biological Sciences 283, 20161304 (2016).

56. I. A. Schneider-Crease et al., Stronger maternal social bonds and higher rank are associated with accelerated infant maturation in Kinda baboons. Animal Behaviour 189, 47–57 (2022).

57. L. Luevano, C. Sutherland, S. J. Gonzalez, R. Hernández-Pacheco, Rhesus macaques compensate for reproductive delay following ecological adversity early in life. Ecology and Evolution 12, e8456 (2022).

58. M. N. Zipple, E. K. Roberts, S. C. Alberts, J. C. Beehner, Male-mediated prenatal loss: Functions and mechanisms. Evolutionary Anthropology: Issues, News, and Reviews 28, 114–125 (2019).

59. E. J. Levy et al., Early life drought predicts components of adult body size in wild female baboons. American journal of biological anthropology 182, 357–371 (2023).

60. E. C. Lange et al., Early life adversity and adult social relationships have independent effects on survival in a wild primate. Science Advances 9, eade7172 (2023).

61. J. Tung, E. A. Archie, J. Altmann, S. C. Alberts, Cumulative early life adversity predicts longevity in wild baboons. Nature communications 7, 1–7 (2016).

62. M. N. Zipple et al., Maternal death and offspring fitness in multiple wild primates. Proceedings of the National Academy of Sciences 118 (2021).

63. G. E. Blomquist, Trade-off between age of first reproduction and survival in a female primate. Biology letters 5, 339–342 (2009).

64. F. B. Bercovitch, J. D. Berard, Life history costs and consequences of rapid reproductive maturation in female rhesus macaques. Behavioral Ecology and Sociobiology 32, 103–109 (1993).

65. R. G. Rawlins, M. J. Kessler, Climate and seasonal reproduction in the Cayo Santiago macaques. American Journal of Primatology 9, 87–99 (1985).

66. S. K. Patterson et al., Early life adversity has sex-dependent effects on survival across the lifespan in rhesus macaques. Philosophical Transactions of the Royal Society B: Biological Sciences 379, 20220456 (2024).

67. J. P. Mersky, C. E. Janczewski, J. Topitzes, Rethinking the measurement of adversity: Moving toward second-generation research on adverse childhood experiences. Child maltreatment 22, 58–68 (2017).

68. Y. Benjamini, Y. Hochberg, Controlling the false discovery rate: a practical and powerful approach to multiple testing. Journal of the royal statistical society. Series B (Methodological), 289–300 (1995).

69. A. F. Richard, R. E. Dewar, M. Schwartz, J. Ratsirarson, Mass change, environmental variability and female fertility in wild Propithecus verreauxi. Journal of Human Evolution 39, 381–391 (2000).

70. K. L. Henrichs et al., Early menarche and childhood adversities in a nationally representative sample. International journal of pediatric endocrinology 2014, 1–8 (2014).

71. R. Boynton-Jarrett, E. W. Harville, A prospective study of childhood social hardships and age at menarche. Annals of epidemiology 22, 731–737 (2012).

72. J. M. Vigil, D. C. Geary, J. Byrd-Craven, A life history assessment of early childhood sexual abuse in women. Developmental Psychology 41, 553 (2005).

73. L. S. Zabin, M. R. Emerson, D. L. Rowland, Childhood sexual abuse and early menarche: the direction of their relationship and its implications. Journal of Adolescent Health 36, 393–400 (2005).

74. D. S. Lee, A. V. Ruiz-Lambides, J. P. Higham, Higher offspring mortality with short interbirth intervals in free-ranging rhesus macaques. Proceedings of the National Academy of Sciences 116, 6057–6062 (2019).

75. R. M. Sibly, J. H. Brown, Mammal reproductive strategies driven by offspring mortality-size relationships. The American Naturalist 173, E185–E199 (2009).

76. A. J. Wilson, J. M. Pemberton, J. G. Pilkington, T. H. Clutton-Brock, L. Kruuk, Trading offspring size for number in a variable environment: selection on reproductive investment in female Soay sheep. Journal of Animal Ecology, 354–364 (2009).

77. D. Gardner, S. Ozanne, K. Sinclair, Effect of the early-life nutritional environment on fecundity and fertility of mammals. Philosophical Transactions of the Royal Society B: Biological Sciences 364, 3419–3427 (2009).

78. K. M. Shreffler, C. N. Joachims, S. Tiemeyer, Maternal Childhood Experiences and Rapid Repeat Pregnancy in a Low-Income, Urban Cohort. Preprint at Research Square 10.21203/rs.3.rs-34758/v1 (2020).

79. A. Wendt, C. M. Gibbs, S. Peters, C. J. Hogue, Impact of increasing inter-pregnancy interval on maternal and infant health. Paediatric and perinatal epidemiology 26, 239–258 (2012).

80. N. Kozuki, N. Walker, Exploring the association between short/long preceding birth intervals and child mortality: using reference birth interval children of the same mother as comparison. BMC public health 13, 1–10 (2013).

81. W. M. Callaghan, M. F. MacDorman, S. A. Rasmussen, C. Qin, E. M. Lackritz, The contribution of preterm birth to infant mortality rates in the United States. Pediatrics 118, 1566–1573 (2006).

82. S. F. Stringham, Grizzly bear reproductive rate relative to body size. Bears: Their Biology and Management, 433–443 (1990).

83. L. Wauters, A. A. Dhondt, Lifetime reproductive success and its correlates in female Eurasian red squirrels. Oikos, 402–410 (1995).

84. B. Thierry, Unity in diversity: lessons from macaque societies. Evolutionary Anthropology: Issues, News, and Reviews 16, 224–238 (2007).

85. B. Marriott, J. Roemer, C. Sultana, An overview of the food intake patterns of the Cayo Santiago rhesus monkeys (Macaca mulatta): report of a pilot study. P R Health Sci J 8, 87–94 (1989).

86. T. Uller, C. Isaksson, M. Olsson, Immune challenge reduces reproductive output and growth in a lizard. Functional Ecology, 873–879 (2006).

87. M. Nystrand, D. Dowling, Effects of immune challenge on expression of life-history and immune trait expression in sexually reproducing metazoans—a meta-analysis. BMC Biol 18, 1–17 (2020).

88. T. H. Clutton-Brock, S. D. Albon, F. E. Guinness, Parental investment and sex differences in juvenile mortality in birds and mammals. Nature 313, 131–133 (1985).

89. L. Kruuk, T. Clutton-Brock, K. Rose, F. Guinness, Early determinants of lifetime reproductive success differ between the sexes in red deer. Proceedings of the Royal Society of London. Series B: Biological Sciences 266, 1655–1661 (1999).

90. E. D. Drake et al., Sex-specific effects of early-life adversity on adult fitness in a wild mammal. Proceedings B 292, 20250192 (2025).

91. S. J. Gonzalez, A. J. Sherer, R. Hernández-Pacheco, Differential effects of early life adversity on male and female rhesus macaque lifespan. Ecology and Evolution 13, e10689 (2023).

92. B. Sadoughi et al., Age and early life adversity shape heterogeneity of the epigenome across tissues in macaques. bioRxiv, 2025.2007. 2013.664445 (2025).

93. R. G. Rawlins, M. J. Kessler, The history of the Cayo Santiago colony. The Cayo Santiago Macaques, 13–45 (1986).

94. A. Widdig et al., Low incidence of inbreeding in a long-lived primate population isolated for 75 years. Behavioral Ecology and Sociobiology 71, 18 (2017).

95. J. G. Vandenbergh, S. Vessey, Seasonal breeding of free-ranging rhesus monkeys and related ecological factors. Reproduction 15, 71–79 (1968).

96. R. Hernandez-Pacheco et al., Managing the Cayo Santiago rhesus macaque population: the role of density. American journal of primatology 78, 167–181 (2016).

97. C. M. Turcotte et al., Quantifying the relationship between bone and soft tissue measures within the rhesus macaques of Cayo Santiago. American Journal of Biological Anthropology 184, e24920 (2024).

98. D. Maestripieri, C. L. Hoffman, “Behavior and social dynamics of rhesus macaques on Cayo Santiago” in Bones, genetics, and behavior of rhesus macaques: Macaca mulatta of Cayo Santiago and beyond. (Springer, 2011), pp. 247–262.

99. D. Langos, L. Kulik, A. Ruiz-Lambides, A. Widdig, Does male care, provided to immature individuals, influence immature fitness in rhesus macaques? PLoS One 10, e0137841 (2015).

100. D. S. Lee, T. M. Mandalaywala, C. Dubuc, A. Widdig, J. P. Higham, Higher early life mortality with lower infant body mass in a free-ranging primate. Journal of Animal Ecology 89, 2300–2310 (2020).

101. M. N. Zipple, E. A. Archie, J. Tung, J. Altmann, S. C. Alberts, Intergenerational effects of early adversity on survival in wild baboons. Elife 8, e47433 (2019).

102. S. K. Patterson et al., Natural animal populations as model systems for understanding early life adversity effects on aging. Integrative and comparative biology 63, 681–692 (2023).

103. K. J. Verhoeven, K. L. Simonsen, L. M. McIntyre, Implementing false discovery rate control: increasing your power. Oikos 108, 643–647 (2005).

