## Supplementary text, figures, and tables for "Early life adversity shapes life history trade-offs between growth and reproduction in free-ranging rhesus macaques"

**This PDF file includes:**

SI text
Figures S1 to S5
Tables S1 to S4
SI References

**Supporting Information Text**

**Supplementary Methods**

Body size measures

All body size measurements were taken following (1) and (2). Trunk length was measured from vertex to the most caudal point on the buttocks, excluding the ischial tuberosities. Total arm length was calculated as the sum of upper and lower arm length. Upper arm length was measured on the extended arm as the distance from the proximal humeral head (top round part of the shoulder) to the center-point of the most lateral bony protuberance on the elbow, and lower arm length was measured on the flexed arm as the distance from the most proximal bony point on the elbow (olecranon) to the most distal bony protrusion above the wrist on the side of the 5th ray (styloid process of ulna). Total leg length was calculated as the sum of upper leg and lateral lower leg lengths. Upper leg length was measured on the lower limb flexed at both the hip and knee as the distance from the lateral-most bony point at the hip joint (trochanterion summum) along the lateral side of the limb to the lateral-most extension of the knee (femorale on the lateral condyle of the femur), and lateral lower leg length was measured from the lateral tibial condyle to the lateral bulge at the ankle (lateral malleolus). All measurements were taken with a tape measure and recorded in centimeters.

Measuring early life adversity

We quantified seven well-established sources of ELA which have been linked to survival in either this or other non-humane primate populations (3-6).

1. Maternal loss: The death of a mother within the first 4 years of life. The loss of a mother significantly increases offspring mortality during lactation and after weaning (7, 8). Maternal loss can occur through natural causes, such as disease or injury, or permanent removal from the population. Although a natural maternal death likely constitutes a greater source of adversity than maternal removal (as it is likely joined by a period of maternal decline), the sudden loss of a healthy mother may still disrupt offspring social and nutritional support. Due to the severe and lasting consequences of both types of loss, we group them together in our analyses.
2. Primiparity: Offspring born to first-time mothers experience greater mortality (9), potentially due to reduced social and/or nutritional support compared to experienced mothers (10).
3. Maternal kin network: In cercopithecine primates, including this species, familial support is important for survival (11). Here, we use maternal kin network size as a proxy for familial support, as both males and females are more likely to interact with maternal kin than other groupmates (12, 13). We measure maternal kin network size as the number of adult females with a relatedness coefficient $\geq 0.063$ (a level at which kin recognition via vocalizations has been documented in this population) present in the group at the time of an individual's birth (14).
4. Matrilineal rank: In rhesus macaques, female dominance hierarchies are typically organized by matrilines composed of closely related female kin. All members of a single matriline generally outrank or underrank all members of any other matriline within the group. This group-level social structure impacts females' access to resources (15), which in turn has consequences for offspring growth and development (16-18). CPRC researchers observe dyadic interactions and assign a matrilineal rank of high, medium, or low to each matriline within each group annually. In instances where matrilineal rank was not available for an individual's birth year, the stability of matriline ranks over time (19) allowed us to infer rank from adjacent years, following the approach described in (6).

5. Competing sibling: Short interbirth intervals (IBIs) are associated with increased offspring mortality in this species (20). Individuals with a younger sibling born within 355 days of their own birth– representing the bottom quartile of IBIs in this population– were considered to have a close-in-age competing sibling (6).
6. Group size: Large group sizes contribute to feeding competition in wild primates (21). We measured early life group size as the number of adult male and females in the group on the day of an individual's birth.
7. Hurricane: Puerto Rico is vulnerable to high-intensity hurricanes, which can lead to widespread environmental and infrastructural damage. For example, following Hurricane Maria in 2017, green vegetation on Cayo Santiago decreased by 63% contributing to significantly hotter temperatures on the island (22). We considered an individual to have been exposed to this adversity if during their first year of life they experienced any of the 3 major hurricanes to have impacted Cayo Santiago in the recent past: Hurricane Hugo (September 18, 1989), Hurricane Georges (September 21, 1998), and Hurricane Maria (September 20, 2017).

For maternal loss, competing sibling, primiparity, and hurricane exposure, we recorded a binary measure: presence=1, absence=0. For group size and kin network size, we standardized these continuous measures based on the mean and standard deviation of these variables within our dataset. For matrilineal rank, we assigned 0, 0.5 and 1 to high, medium, and low matrilineal rank, respectively. To quantify cumulative ELA, we scaled each continuous ELA variable between 0 and 1, with 1 indicating the highest degree of adversity. We then summed all ELA sources together to generate a composite metric, a method which has been effectively used in previous studies to predict proxies of fitness in both humans and non-human primates (3, 6, 23, 24). Although cumulative ELA in theory could've ranged from 0 - 7, in our dataset it ranged from 0.41 - 4.69. For the 2,100 females in our dataset, we have complete ELA data (all 7 ELA sources + cumulative ELA) for 1,640 individuals.

**Figures**

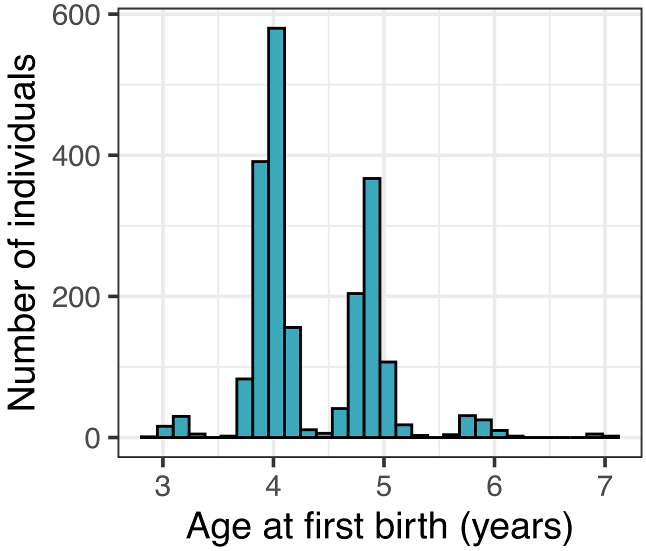


**Fig. S1.** Distribution of age at first birth in the Cayo Santiago macaques. Seasonal breeding accounts for the modal distribution of ages (mean= 4.3, range=2.9-7.1 years).

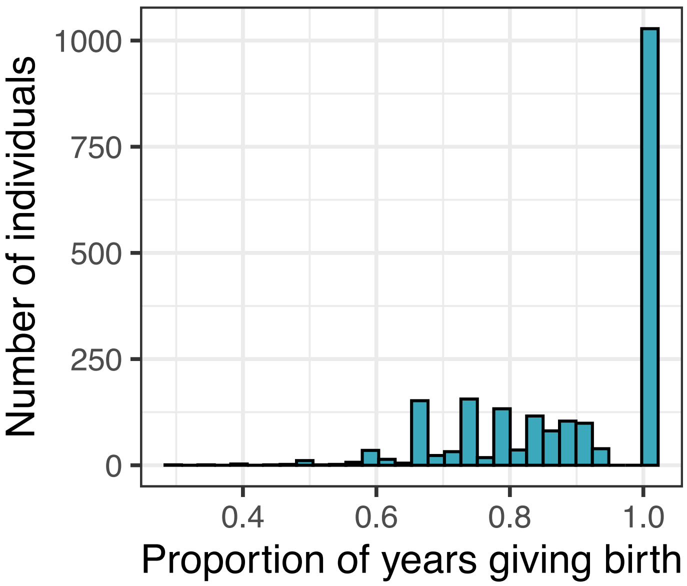


**Fig. S2.** Distribution of proportion of birth seasons in which a female gives birth (mean=0.89, range=0.3-1).

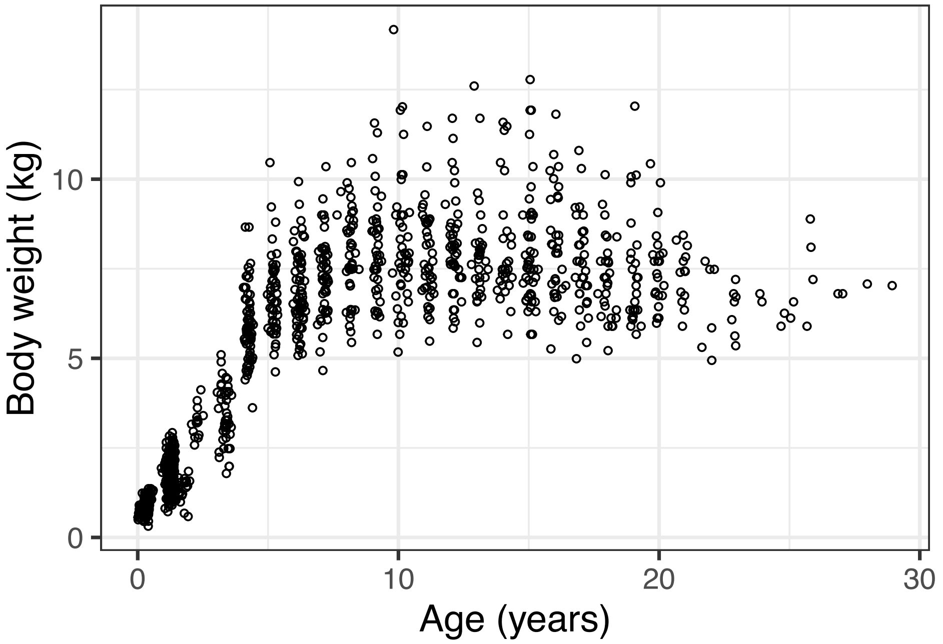


**Fig. S3.** Body weight measurements across the life course.

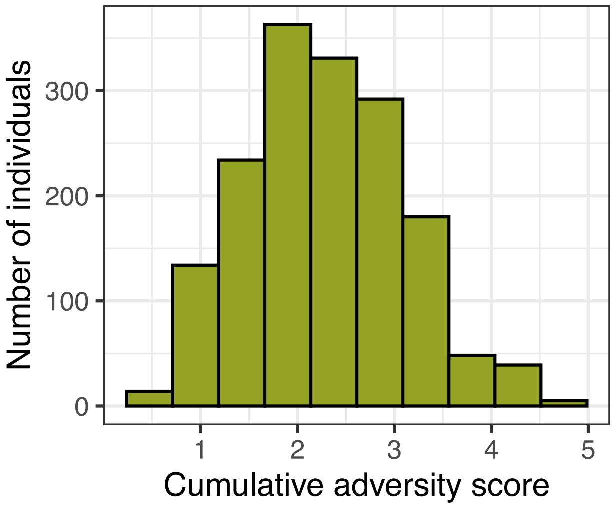


**Fig S4.** Distribution of cumulative ELA scores among individuals included in our analyses.

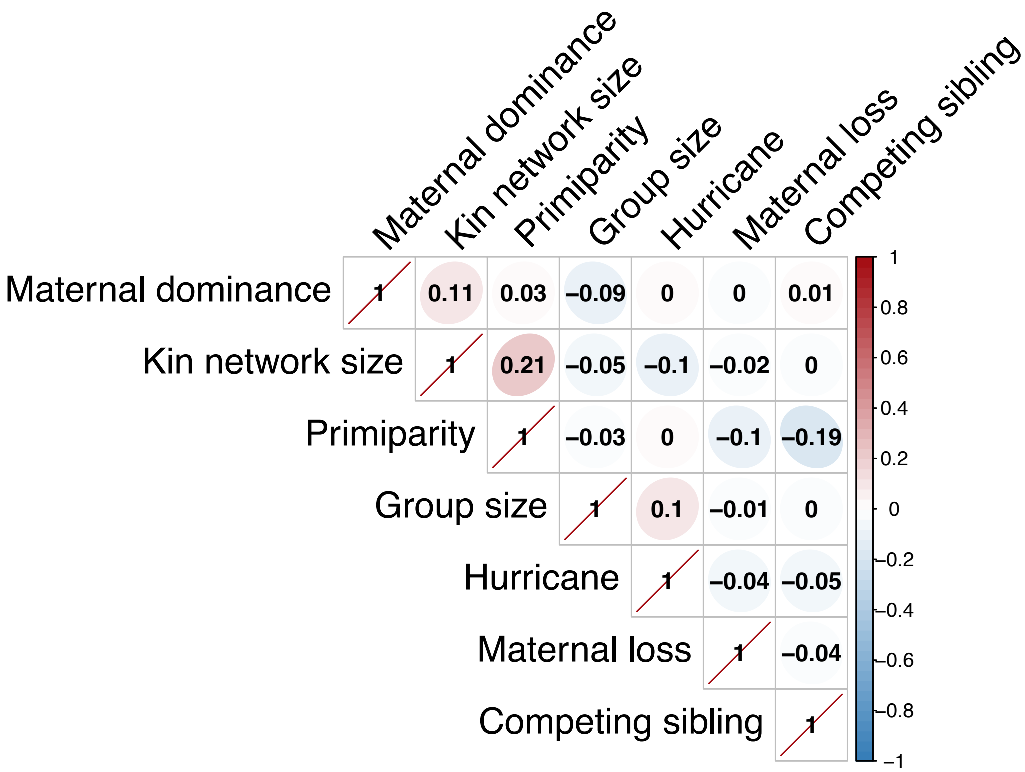


**Fig S5.** Correlation matrix demonstrating the lack of correlated exposure to different individual sources of adversity.

**Tables**

**Table S1.** Summary of early-life adversity metrics. Definitions of each adversity as well as the data type, number of individuals assessed for the adversity and the number of individuals who experienced the adversity for non-continuous metrics. Numbers are presented for the reproduction and growth datasets separately.

| Type of adversity | Definition | Data type | Number measured (reproduction;growth) | Number experienced (reproduction;growth) |
|---|---|---|---|---|
| Maternal loss | Death of mother before reaching 4 years old | Binary | 2100;1327 | 408;351 |
| Maternal primiparity | First born offspring to a mother | Binary | 2100;1327 | 391;213 |
| Competing sibling | Birth of a younger sibling <355 days after date of birth | Binary | 2100;1327 | 423;266 |
| Kin network size | Number of closely related adult female relatives on date of birth | Continuous | 2091;1323 | n/a |
| Group size | Number of adults in the group on date of birth | Continuous | 2098;1327 | n/a |
| Maternal dominance | Categorical dominance (high/medium/low) of the mother's matriline in year of birth | Ternary | 1649;1105 | Mid ranking: 592;352<br>Low ranking: 646;582 |
| Hurricane | Experienced a major hurricane during 1st year of life | Binary | 2100;1327 | 115;34 |

**Table S2.** Model results assessing the relationship between ELA and life history. Each line represents the results of a separate model.

| Response | Predictor | n | Estimate | Std. Error | p-value | FDR |
|---|---|---|---|---|---|---|
| Age at first birth | Maternal Loss | 2100 | 0.0680629 | 0.02950005 | 0.02114138 | 0.0475681 |
| Age at first birth | Primiparity | 2100 | 0.01512298 | 0.02993625 | 0.61349129 | 0.6901777 |
| Age at first birth | Competing Sibling | 2100 | 0.00383402 | 0.02942823 | 0.89635452 | 0.89635452 |
| Age at first birth | Group size | 2098 | 0.07326212 | 0.01757497 | 3.73E-05 | 0.00016804 |
| Age at first birth | Kin network size | 2091 | -0.0060054 | 0.01020906 | 0.55643693 | 0.6901777 |
| Age at first birth | Matriline rank- mid v high | 1649 | 0.07236556 | 0.03622747 | 0.04612828 | 0.0830309 |
| Age at first birth | Hurricane | 2100 | 0.09897353 | 0.11518144 | 0.39341746 | 0.59012619 |
| Age at first birth | Cumulative Adversity | 1640 | 0.06443009 | 0.01762419 | 0.00026609 | 0.00079826 |
| Age at first birth | Matriline rank- low v high | 1649 | 0.15933678 | 0.03546434 | 7.89E-06 | 7.11E-05 |
| Reproductive rate | Maternal Loss | 1419 | -0.1043724 | 0.07380189 | 0.15729619 | 0.28313315 |
| Reproductive rate | Primiparity | 1419 | -0.0588594 | 0.07572814 | 0.43701379 | 0.56187488 |
| Reproductive rate | Competing Sibling | 1419 | 0.21519929 | 0.07875177 | 0.00628313 | 0.03411185 |
| Reproductive rate | Group size | 1417 | 0.10550382 | 0.03982796 | 0.00807331 | 0.03411185 |
| Reproductive rate | Kin network size | 1411 | -0.0099976 | 0.0252231 | 0.69183426 | 0.72098131 |
| Reproductive rate | Matriline rank- mid v high | 1291 | -0.2201211 | 0.08696666 | 0.01137062 | 0.03411185 |
| Reproductive rate | Hurricane | 1419 | 0.19840825 | 0.16701539 | 0.23484766 | 0.3522715 |
| Reproductive rate | Cumulative Adversity | 1282 | -0.0155268 | 0.04347455 | 0.72098131 | 0.72098131 |
| Reproductive rate | Matriline rank- low v high | 1291 | -0.1918618 | 0.0937259 | 0.04065296 | 0.09146917 |
| Sub-adult weight | Maternal Loss * age | 612 | -0.0577917 | 0.03662488 | 0.11510957 | 0.15347943 |
| Sub-adult weight | Primiparity * age | 612 | 0.0180168 | 0.0498963 | 0.71816255 | 0.72105682 |
| Sub-adult weight | Competing sibling * age | 612 | -0.1706815 | 0.04043663 | 2.81E-05 | 0.00022493 |
| Sub-adult weight | Group size * age | 612 | 0.08111507 | 0.02013653 | 6.36E-05 | 0.00025454 |
| Sub-adult weight | kin network size * age | 610 | 0.02774195 | 0.01525053 | 0.06939939 | 0.11132909 |
| Sub-adult weight | Cumulative Adversity * age | 487 | -0.0467663 | 0.02317534 | 0.04416418 | 0.11132909 |
| Sub-adult weight | Matriline rank- mid v high * age | 488 | 0.12233054 | 0.06726106 | 0.06958068 | 0.11132909 |
| Sub-adult weight | Matriline rank- low v high * age | 488 | -0.0219305 | 0.06137927 | 0.72105682 | 0.72105682 |
| Adult weight | Maternal Loss | 697 | -0.0082632 | 0.00711114 | 0.24594672 | 0.43055055 |
| Adult weight | Primiparity | 697 | 0.00461897 | 0.00830977 | 0.57863803 | 0.65096778 |
| Adult weight | Competing Sibling | 697 | 0.00813272 | 0.00725377 | 0.26292795 | 0.43055055 |
| Adult weight | Group size | 697 | -0.0078157 | 0.00728459 | 0.2870337 | 0.43055055 |
| Adult weight | Kin network size | 695 | -0.0031265 | 0.00246099 | 0.20468264 | 0.43055055 |
| Adult weight | Matriline rank- mid v high | 605 | -0.018017 | 0.01138748 | 0.11474722 | 0.43055055 |
| Adult weight | Hurricane | 697 | 0.01255501 | 0.02937345 | 0.6731346 | 0.6731346 |
| Adult weight | Cumulative Adversity | 603 | -0.0036495 | 0.00460724 | 0.42887332 | 0.55140856 |
| Adult weight | Matriline rank- low v high | 605 | -0.0325143 | 0.01239569 | 0.00937637 | 0.08438735 |
| Adult arm length | Maternal Loss | 393 | -0.0022487 | 0.00311766 | 0.47138908 | 0.7870597 |
| Adult arm length | Primiparity | 393 | -0.0033198 | 0.00343259 | 0.33437583 | 0.75234561 |
| Adult arm length | Competing Sibling | 393 | 0.00045095 | 0.00321102 | 0.88842387 | 0.93741704 |
| Adult arm length | Group size | 393 | 0.00295928 | 0.00255889 | 0.2487257 | 0.75234561 |

| | | | | | | |
|---|---|---|---|---|---|---|
| Adult arm length | Kin network size | 393 | 9.19E-05 | 0.00116923 | 0.93741704 | 0.93741704 |
| Adult arm length | Matriline rank- mid v high | 311 | 0.00284961 | 0.00504471 | 0.57374164 | 0.7870597 |
| Adult arm length | Hurricane | 393 | -0.0135146 | 0.00788617 | 0.08790384 | 0.75234561 |
| Adult arm length | Cumulative Adversity | 311 | -0.0022095 | 0.00207211 | 0.28800151 | 0.75234561 |
| Adult arm length | Matriline rank- low v high | 311 | 0.00275399 | 0.00533024 | 0.61215755 | 0.7870597 |
| Adult leg length | Maternal Loss | 389 | -0.0043943 | 0.00313511 | 0.16228071 | 0.67572707 |
| Adult leg length | Primiparity | 389 | -0.0014282 | 0.00344853 | 0.67913012 | 0.89438641 |
| Adult leg length | Competing Sibling | 389 | -0.0003026 | 0.0032294 | 0.92543304 | 0.92543304 |
| Adult leg length | Group size | 389 | -0.0011273 | 0.00345917 | 0.74495812 | 0.89438641 |
| Adult leg length | Kin network size | 389 | -0.0006196 | 0.00117626 | 0.59883933 | 0.89438641 |
| Adult leg length | Matriline rank- mid v high | 307 | 0.00135176 | 0.00519237 | 0.79501014 | 0.89438641 |
| Adult leg length | Hurricane | 389 | -0.0106124 | 0.00804183 | 0.18837467 | 0.67572707 |
| Adult leg length | Cumulative Adversity | 307 | -0.0025349 | 0.00208227 | 0.22524236 | 0.67572707 |
| Adult leg length | Matriline rank- low v high | 307 | 0.00258318 | 0.00608542 | 0.67251583 | 0.89438641 |
| Adult trunk length | Maternal Loss | 390 | -0.0042645 | 0.00250171 | 0.0895217 | 0.29647135 |
| Adult trunk length | Primiparity | 390 | -0.0013536 | 0.00273085 | 0.62062991 | 0.7890289 |
| Adult trunk length | Competing Sibling | 390 | 0.00310238 | 0.00258167 | 0.23069912 | 0.41525842 |
| Adult trunk length | Group size | 390 | 0.0011568 | 0.00301242 | 0.70135903 | 0.7890289 |
| Adult trunk length | Kin network size | 390 | -0.0011366 | 0.00093153 | 0.2237576 | 0.41525842 |
| Adult trunk length | Matriline rank- mid v high | 310 | 0.0004659 | 0.00390384 | 0.90520151 | 0.90520151 |
| Adult trunk length | Hurricane | 390 | -0.0137035 | 0.00664808 | 0.04052355 | 0.29647135 |
| Adult trunk length | Cumulative Adversity | 310 | -0.0027008 | 0.00162656 | 0.09882378 | 0.29647135 |
| Adult trunk length | Matriline rank- low v high | 310 | -0.0039039 | 0.00486347 | 0.42357498 | 0.63536246 |

**Table S3.** Model results assessing the relationship between life history domains. Each line represents the results of a separate model.

| Response | Predictor | n | Estimate | Std. Error | p-value | FDR |
|---|---|---|---|---|---|---|
| Age at first birth | sub-adult weight | 203 | -0.3108013 | 0.10079318 | 0.004185 | n/a |
| Age at first birth | Maternal loss * sub-adult weight | 201 | 0.08904159 | 0.21407548 | 0.67791162 | 0.77475613 |
| Age at first birth | Primiparity * sub-adult weight | 201 | 0.0560447 | 0.21541368 | 0.79500279 | 0.79500279 |
| Age at first birth | Competing sibling * sub-adult weight | 201 | -0.1316742 | 0.28689566 | 0.64676787 | 0.77475613 |
| Age at first birth | Group weight * sub-adult weight | 201 | -0.3011952 | 0.14436404 | 0.0382553 | 0.30604241 |
| Age at first birth | kin network weight * sub-adult weight | 201 | -0.0623878 | 0.11598671 | 0.59126213 | 0.77475613 |
| Age at first birth | Cumulative adversity * sub-adult weight | 163 | 0.07066575 | 0.14652833 | 0.63030462 | 0.77475613 |
| Age at first birth | Matriline rank- mid v high * sub-adult weight | 163 | 0.38853802 | 0.33577041 | 0.2490209 | 0.66652113 |
| Age at first birth | Matriline rank- low v high * sub-adult weight | 163 | 0.36021328 | 0.3119089 | 0.24994543 | 0.66652113 |
| Reproductive rate | Adult weight | 353 | 1.54715717 | 0.68217771 | 0.02333116 | n/a |
| Reproductive rate | Maternal loss * adult weight | 353 | 3.41573456 | 1.69096741 | 0.04338459 | 0.08676918 |
| Reproductive rate | Primiparity * adult weight | 353 | -5.6842098 | 1.95903728 | 0.00371343 | 0.02970741 |
| Reproductive rate | Competing sibling * adult weight | 353 | -3.5493325 | 1.66935421 | 0.03348903 | 0.08676918 |
| Reproductive rate | Group weight * adult weight | 353 | 0.65613069 | 0.66817223 | 0.3261105 | 0.37269771 |
| Reproductive rate | kin network weight * adult weight | 352 | -0.1895633 | 0.37831179 | 0.61631705 | 0.61631705 |
| Reproductive rate | Cumulative adversity * adult weight | 298 | -0.9901549 | 0.95176849 | 0.29818585 | 0.37269771 |
| Reproductive rate | Matriline rank- mid v high * adult weight | 299 | 3.90996284 | 1.85199101 | 0.0347533 | 0.08676918 |
| Reproductive rate | Matriline rank- low v high * adult weight | 163 | 2.49311888 | 1.82081291 | 0.17092615 | 0.27348184 |

**Table S4.** Model results assessing the relationship between life history and reproductive success. We fit a separate model for each life history trait, including age at first birth and body weight as linear terms and proportion of seasons giving birth (reproductive rate) as both linear and quadratic.

| Response | Predictor | n | Estimate | Std. Error | p-value |
|---|---|---|---|---|---|
| Number of surviving offspring | Age at first birth | 279 | -0.6573002 | 0.27063898 | 0.0158109 |
| Number of surviving offspring | Reproductive rate^2 | 279 | -57.983896 | 10.2592745 | 3.97E-08 |
| Number of surviving offspring | Body weight | 52 | -1.0234139 | 0.54258906 | 0.06508777 |

### SI References


1. A. Schultz, "The technique of measuring the outer body of human fetuses and primates in general" in Contributions to embryology. (Carnegie Institution of Washington, 1929), vol. 117, pp. 213-257.
2. S. C. Antón *et al.*, Integrative measurement protocol for morphological and behavioral research in human and nonhuman primates. *Am J Phys Anthropol S* **48**, 78-79 (2009).
3. J. Tung, E. A. Archie, J. Altmann, S. C. Alberts, Cumulative early life adversity predicts longevity in wild baboons. *Nature communications* **7**, 1-7 (2016).
4. C. J. Weibel, J. Tung, S. C. Alberts, E. A. Archie, Accelerated reproduction is not an adaptive response to early-life adversity in wild baboons. *Proceedings of the National Academy of Sciences* **117**, 24909-24919 (2020).
5. M. N. Zipple, E. A. Archie, J. Tung, J. Altmann, S. C. Alberts, Intergenerational effects of early adversity on survival in wild baboons. *Elife* **8**, e47433 (2019).
6. S. K. Patterson *et al.*, Early life adversity has sex-dependent effects on survival across the lifespan in rhesus macaques. *Philosophical Transactions of the Royal Society B: Biological Sciences* **379**, 20220456 (2024).
7. M. A. Stanton, E. V. Lonsdorf, C. M. Murray, A. E. Pusey, Consequences of maternal loss before and after weaning in male and female wild chimpanzees. *Behavioral Ecology & Sociobiology* **74**, 1-11 (2020).
8. M. N. Zipple *et al.*, Maternal death and offspring fitness in multiple wild primates. *Proceedings of the National Academy of Sciences* **118** (2021).
9. F. B. Bercovitch, M. R. Lebron, H. S. Martinez, M. J. Kessler, Primigravidity, body weight, and costs of rearing first offspring in rhesus macaques. *American Journal of Primatology* **46**, 135-144 (1998).
10. F. Pittet, K. Hinde, Meager Milk: Lasting consequences for adult daughters of primiparous mothers among rhesus macaques (Macaca Mulatta). *Integrative and comparative biology* **63**, 569-584 (2023).
11. L. J. N. Brent, A. Ruiz-Lambides, M. L. Platt, Family network size and survival across the lifespan of female macaques. *Proceedings of the Royal Society B: Biological Sciences* **284** (2017).
12. L. Kulik, F. Amici, D. Langos, A. Widdig, Sex differences in the development of social relationships in rhesus macaques (Macaca mulatta). *International Journal of Primatology* **36**, 353-376 (2015).
13. A. Widdig, D. Langos, L. Kulik, Sex differences in kin bias at maturation: male rhesus macaques prefer paternal kin prior to natal dispersal. *American Journal of Primatology* **78**, 78-91 (2016).
14. D. Rendall, P. S. Rodman, R. E. Emond, Vocal recognition of individuals and kin in free-ranging rhesus monkeys. *Animal behaviour* **51**, 1007-1015 (1996).
15. M. L. Boccia, M. Laudenslager, M. Reite, Food distribution, dominance, and aggressive behaviors in bonnet macaques. *American Journal of Primatology* **16**, 123-130 (1988).
16. C. Garcia, P. C. Lee, L. Rosetta, Growth in colony living anubis baboon infants and its relationship with maternal activity budgets and reproductive status. *American Journal of Physical Anthropology* **138**, 123-135 (2009).
17. S. E. Johnson, Life history and the competitive environment: trajectories of growth, maturation, and reproductive output among chacma baboons. *American Journal of Physical Anthropology* **120**, 83-98 (2003).
18. I. A. Schneider-Crease *et al.*, Stronger maternal social bonds and higher rank are associated with accelerated infant maturation in Kinda baboons. *Animal Behaviour* **189**, 47-57 (2022).
19. B. Thierry, Unity in diversity: lessons from macaque societies. *Evolutionary Anthropology: Issues, News, and Reviews* **16**, 224-238 (2007).

20. D. S. Lee, A. V. Ruiz-Lambides, J. P. Higham, Higher offspring mortality with short interbirth intervals in free-ranging rhesus macaques. *Proceedings of the National Academy of Sciences* **116**, 6057-6062 (2019).
21. C. P. van Schaik, M. A. van Noordwijk, R. J. de Boer, I. den Tonkelaar, The effect of group size on time budgets and social behaviour in wild long-tailed macaques (Macaca fascicularis). *Behavioral Ecology & Sociobiology* **13**, 173-181 (1983).
22. C. Testard *et al.*, Ecological disturbance alters the adaptive benefits of social ties. *Science* **384**, 1330-1335 (2024).
23. E. A. Schilling, R. H. Aseltine, S. Gore, The impact of cumulative childhood adversity on young adult mental health: Measures, models, and interpretations. *Social Science & Medicine* **66**, 1140-1151 (2008).
24. V. J. Felitti *et al.*, Relationship of childhood abuse and household dysfunction to many of the leading causes of death in adults: The Adverse Childhood Experiences (ACE) Study. *American Journal of Preventive Medicine* **14**, 245-258 (1998).
